# IL4i1-sourced oncometabolites promote neuroblastoma cell survival

**DOI:** 10.1101/2025.08.07.669062

**Authors:** Camille Guyot, Lee-Ann Van de Velde, Lisa Kainacher, Johanna Ruoff, Ana Bici, Nicolas Bernard Peterson, E. Kaitlynn Allen, Barbara Steigenberger, Assa Yeroslaviz, Jun Yang, Leonie Zeitler, Paul G. Thomas, Peter J. Murray

## Abstract

High-risk neuroblastoma (NB) is driven by the amplification of MYCN in conjunction with additional oncogenic mutations in kinases such as ALK. NB cells require antioxidant responses to maintain redox balance and are highly sensitive to ferroptosis. Here, we show that metabolites derived from infiltrating immune cells expressing IL4i1, a secreted oxidoreductase, are potent suppressors of NB ferroptosis. IL4i1 metabolites (indole-3-pyruvate and 4-hydroxyphenylpyruvate) blocked ferroptosis in all human NB cell lines via a mechanism that depended on free radical scavenging and NRF2 activation but did not require the aryl hydrocarbon receptor. Supernatant transfer experiments confirmed that IL4i1 creates a milieu that protects NB cells from oxidative cell death. Importantly, mice lacking IL4i1 were protected from NB in a high-penetrance MYCN and mutant ALK-driven autochthonous cancer model. Therefore, we propose that immune IL4i1 is permissive for NB growth and survival. IL4i1 produces context-dependent oncometabolites and, as a secreted enzyme, represents a target for cell death manipulation in cancers sensitive to oxidative stress-driven cell death.

**Graphical abstract:** 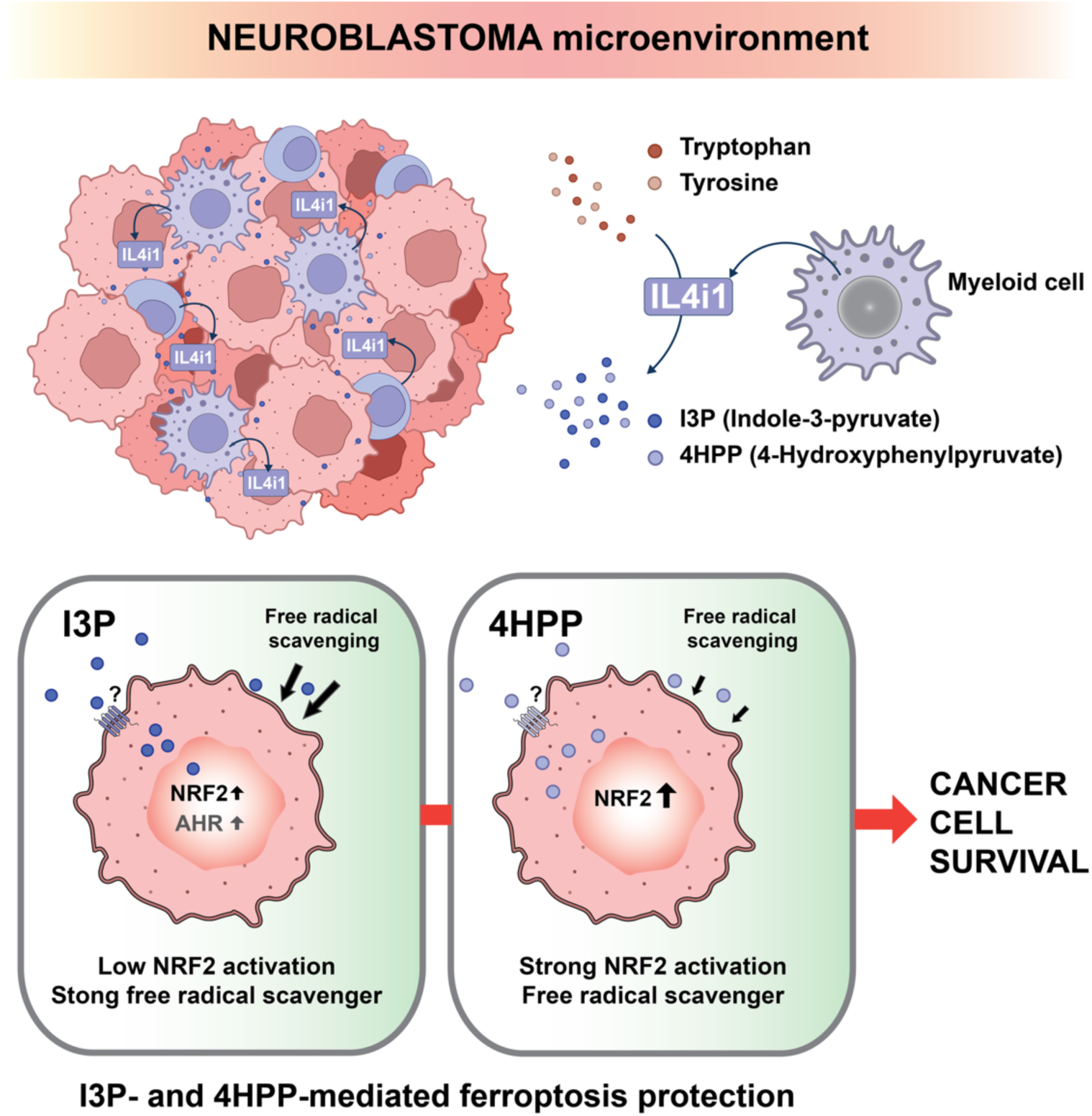

## Introduction

Hallmarks of high-risk neuroblastoma (NB) include amplification of *MYCN* ^1^, activating mutations in the ALK tyrosine kinase receptor (encoded by *ALK*), and a suite of downstream transcriptional, epigenetic and proteomic changes linked to MYCN-driven tumor growth ^2–4^. NB originates during the development of the peripheral sympathetic nervous system and generally manifests as a malignancy of the adrenal medulla ^5,6^. As the most common childhood solid tumor ^7^, extensive genetic, phenotypic and clinical research has defined the cells-of-origin of high-risk NB, their phenotypes (adrenergic versus mesenchymal) and the roles of driver oncogenes in the malignant phenotype. Furthermore, mouse models that recapitulate many aspects of the disease, along with well-characterized human NB cell lines have advanced the understanding of the genotype-phenotype relationships in different NB risk categories and their association with clinical manifestations of NB ^4,8–16^.

Oncogene-driven proliferation drives an increased demand for maintenance of the cell-intrinsic redox balance, which is mediated by key pathways such as glutathione and CoQ homeostasis ^17–19^. Consistent with MYCN-driven NB proliferation, high-risk NB cells undergo terminal lipid peroxidation and ferroptosis when glutathione amounts are perturbed by cysteine depletion, for example, by interfering with the activity of SLC7A11, the key cystine-glutamate exchange transporter ^20–23^. Furthermore, high-risk NB cells critically depend on the maintenance of iron homeostasis and the transsulfuration pathway (TSP) to withstand disruptions in cellular redox balance ^21–23^ amongst other metabolic rewiring events ^24,25^. Like most rapidly proliferating cells, high-risk NB must maintain sufficient levels of glutathione peroxidase 4 (GPX4) to support the glutathione redox cycle. GPX4 contains a single selenocysteine in its catalytic site and its biosynthesis depends on the machinery that incorporates selenocysteine into nascent proteins ^26^. NB cells acquire selenium via LRP8-mediated uptake of SELENOP ^20^, a serum selenocysteine “sink” protein and perturbation of this pathway is an additional ferroptosis vulnerability of NB. Other key redox enzymes, such as thioredoxin reductase (TXNRD1), which also rely on a single selenocysteine residue for their enzymatic activity, may also be essential for the growth and proliferation of NB cells ^21^. Taken together, high-risk NB cells are vulnerable to disruptions in pathways that regulate cellular redox homeostasis.

Cell-intrinsic pathways to suppress oxidative stress are one arm of the body’s defenses against the effects of excessive free radicals, lipid peroxidation and ferroptotic cell death ^17,27^. A second cell-extrinsic defensive branch encompasses factors including diet-derived antioxidants, such as vitamin E and other plant-derived radical scavengers. However, there is also an endogenous pathway that generates anti-ferroptotic molecules from tryptophan and tyrosine metabolism, which centers on immune cells that express IDO1 and IL4i1 ^28^. IDO1 metabolizes tryptophan to kynurenine, and downstream metabolites from this pathway are potent inhibitors of ferroptosis that act via free radical scavenging (FRS), modulation of cystine importation, NRF2 activation and responses to amino acid starvation ^29,30^. While IDO1 is cytoplasmic, IL4i1 is secreted, predominantly from activated myeloid cells and B cells ^28,29,31,32^. IL4i1 metabolizes aromatic amino acids to their keto acids via oxidative deamination. The IL4i1-derived keto acids from tryptophan and tyrosine metabolism - indole-3-pyruvate (I3P) and 4-hydroxyphenyl-pyruvate (4HPP), respectively - inhibit ferroptosis, whereas phenylpyruvate (PP), derived from phenylalanine, does not ^33^. Similar to many other indoles derived from food and the microbiome, I3P also activates the aryl hydrocarbon receptor (AHR) ^34^. However, the precise mechanisms of how these metabolites modulate networks of cellular oxidative stress responses and the AHR are unclear ^28^. Because IL4i1 generates anti-ferroptotic metabolites outside the cell by virtue of being a secreted enzyme, it can potentially influence oxidative stress responses in neighboring cells through metabolite signaling. Thus, remnant “persister” cells that survive chemotherapy ^19,27,35–37^ or malignant cells with high sensitivity to oxidative stress damage and ferroptosis could harness the IL4i1 pathway as a survival mechanism.

High-risk NBs are populated with immune cells ^6,16,38–41^, and the degree of immune infiltration is correlated with clinical outcome but inversely correlated to the adrenergic-mesenchymal phenotype: thus, *MYCN* amplification in adrenergic NB cells controls immune evasion ^6,38,42^. By contrast, reversible conversion of NB to the mesenchymal state may be involved in chemotherapy resistance ^5,6,42–45^. Conceivably, if one or more immune populations express IL4i1 in high-risk NB, then stress-sensitive tumor cells could harness I3P or 4HPP to survive redox stress beyond their cell-intrinsic metabolic re-wiring: IL4i1 would therefore represent an immune-specific target enzyme for high-risk NB, or indeed any other tumor type dependent on escaping the effects of oxidative stress. Here, we explored the links between ferroptosis in high-risk NB and IL4i1. Our findings indicate that IL4i1-derived metabolites not only suppress ferroptosis in high-risk NB cell lines, but IL4i1 is also a central factor in the development and expansion of NB in a high-penetrance model of amplified MYCN- and mutant ALK-driven NB.

## Results

### IL4i1 expression in human and mouse high-risk NB

Single-cell RNA-seq experiments in human cancers show that different immune cell types in the tumor microenvironment (TME) express IL4i1 mRNA, including B cells, myeloid cells, and especially activated dendritic cells (mregDCs), macrophages or stromal cells ^28,46^. We first examined the specificity of IL4i1 mRNA expression in a harmonized, single-cell/single-nucleus transcriptomic atlas of human neuroblastoma ^47^. As expected, IL4i1 mRNA^+^ cells included lymphocytes and myeloid cells. Importantly, NB tumor cells, defined by their expression of *PHOX2B* ^4^, did not express IL4i1 mRNA (**Figure 1A and Figure S1A**). A concordant pattern to human NB was also observed in scRNAseq and spatial RNAseq data from tumors from the TH-MYCN; *Alk*^F1178L^ mice ^8^, where IL4i1 mRNA expression was detected sparsely throughout the TME consistent with expression in fractions of myeloid and B cells we observed in scRNAseq (**Figure 1B, S1B and S1C).** Similarly, established NB human cell lines, which bear different combinations of amplified MYCN alleles and ALK mutations, did not express IL4i1 protein (**Figure 1C** and **1D**). Therefore, IL4i1 expression in human NB and in mouse model of high-risk NB is restricted to immune cells within the TME.

**Figure 1.**
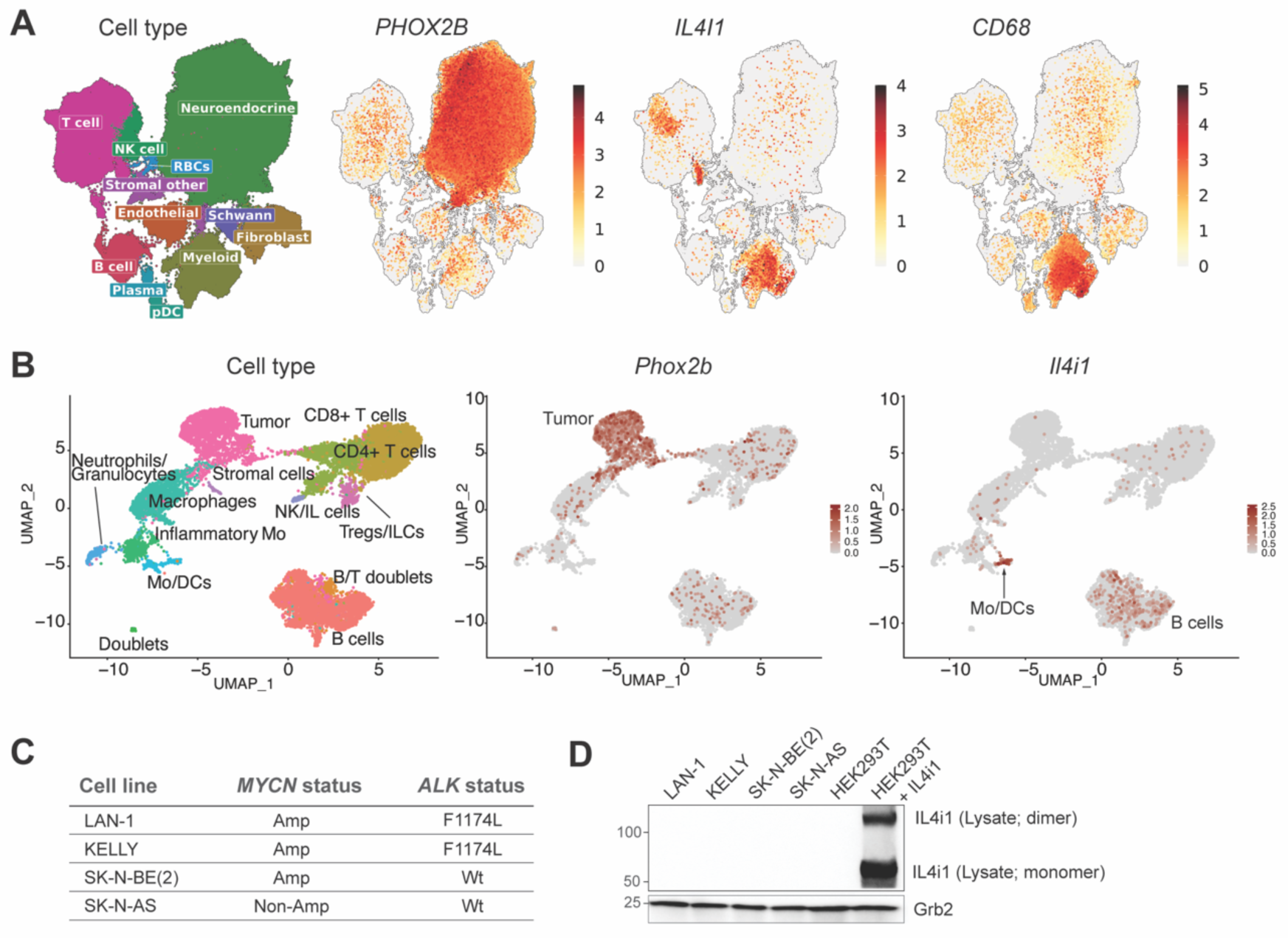
scRNAseq analysis of IL4i1 expression in NB. **(A)** Overview of scRNAseq populations from human NB datasets ^47^. PHOX2B mRNA is enriched in the tumor cells while IL4I1 mRNA is enriched in myeloid and T cells. CD68 serves as a marker for myeloid cells. **(B)** scRNAseq analysis of IL4i1 mRNA^+^ cells in TH-MYCN; *Alk*^F1178L^ NB ^8^. Phox2b mRNA is enriched in the tumor cells while IL4i1 mRNA is enriched in Mo/DCs and B cells. **(C)** Genomic status of *MYCN* and *ALK* in the human NB cell lines. Amp: *MYCN* amplified; Non-Amp: *MYCN* non-amplified; Wt: wild type sequence. **(D)** Lysates of the human NB cell lines were probed with an antibody directed against human IL4i1. HEK293T cells were transiently transfected with a cDNA encoding IL4i1 (IL4i1^WT^) as the positive control for IL4i1 expression.

### IL4i1-derived metabolites protect NB cells from induced ferroptosis

Human NB cells rewire their cysteine, glutathione and selenium metabolism to protect themselves from oxidative stress. However, NB cells within the TME are surrounded by other cells and especially immune cells, which may influence their survival, proliferation and differentiation. As IL4i1 is a secreted enzyme, which metabolizes tryptophan and tyrosine to I3P and 4HPP, respectively, in the extracellular space ^28^, we considered that NB cells undergoing oxidative stress could use IL4i1-derived metabolites to protect themselves from ferroptosis. To test this idea, we used LAN-1 NB cells, which are *MYCN*-amplified and carry an activating mutation in *ALK* (F1174L) ^2^, and challenged them with erastin, which blocks the import of cystine via SLC7A11. We also used RSL3, which inhibits GPX4, TXNRD1 and other selenoproteins and rapidly disrupts cellular redox homeostasis ^48,49^. We measured the response of LAN-1 cells to these redox-corrupting agents using time-resolved live-cell imaging of cell viability. When we added I3P or 4HPP at concentrations of 25 µM and 50 µM, we observed a dose-dependent protection from cell death, with 50 µM providing a level of protection comparable to ferrostatin-1 (Fer-1), a control inhibitor of ferroptosis (**Figure 2A**). Notably, I3P exhibited a greater protective effect than 4HPP (**Figure 2A** and **2B**).

**Figure 2.**
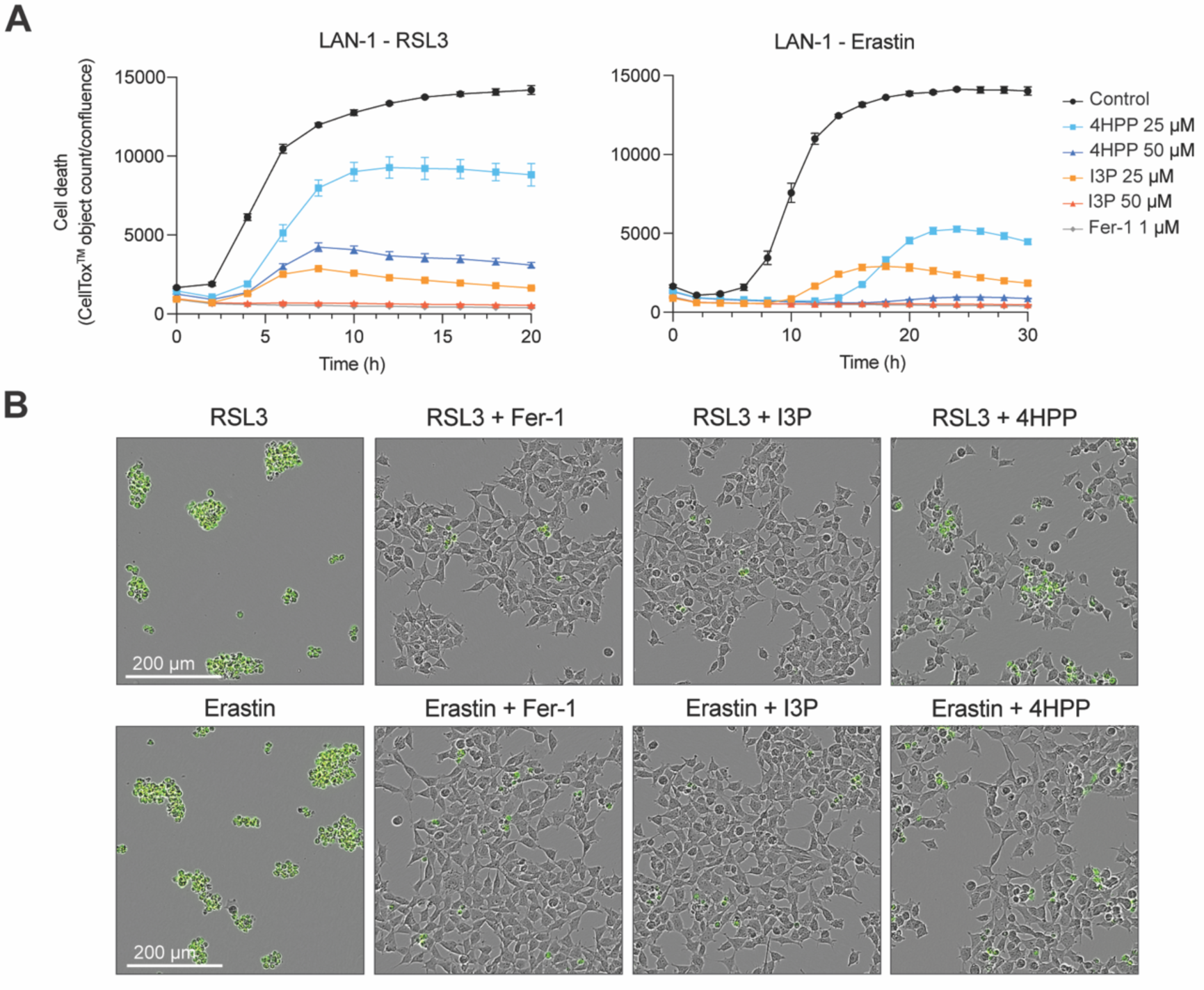
IL4i1 metabolites protect human NB cells from ferroptosis **(A)** LAN-1 cells were challenged with 0.5 µM erastin or 0.1 µM RSL3 in the presence or absence of I3P, 4HPP or Fer-1 at the indicated concentration and cell death monitored by live cell imaging (Incucyte) using CellTox^TM^ Green over time. The control was treated with media. Data are presented as mean ± s.e.m., n = 3 samples. The experiment was replicated 3 times. **(B)** Representative images of LAN-1 cells challenged with RSL3 or erastin for 30 h with or without 50 µM I3P, 4HPP or 1 µM Fer-1.

### Quantification of IL4i1-derived metabolites

To measure the amounts of IL4i1-derived metabolites in tumors, ideally, one would want both time-resolved and spatial information about metabolite production in relation to amino acid consumption. However, when we used untargeted metabolomics on tumors derived from TH-MYCN; *Alk*^F1178L^ mice, we found hundreds of changes relative to the serum of the same mice or mice lacking tumors (**Figure S2A**). While tryptophan was depleted, as expected from previous studies with mouse models of PDAC ^50^, quantification of the IL4i1 metabolites was complicated by the fact that a comparator of a tumor devoid of IL4i1 metabolites was not possible (discussed further below). Further, IL4i1^+^ immune cells make up a small fraction of the total NB tumor (**Figure 1A, 1B and S1C**), which suggests that a single timepoint measurement of IL4i1 metabolites from a complex tumor environment is unlikely to provide definitive information about IL4i1 metabolites or their relationship to tumor cell survival. To reduce the complexity of quantifying the relationship between IL4i1 metabolite concentrations and their biological effects, we tested if IL4i1 itself generated an anti-ferroptotic milieu that would influence the survival of stressed NB cells. We expressed human IL4i1 (IL4i1^WT^) or an enzyme-dead IL4i1 mutant (IL4i1^ED^) ^33^ in HEK293T cells and transferred the unmanipulated supernatant to NB cells (**Figure 3A, S2B and S2C**). Under these conditions, secreted IL4i1^WT^ (or IL4i1^ED^) was also transferred to the recipient cells and remained active. In parallel, we filtered the supernatant to remove all material >3 kDa ^51^, which would retain the metabolic products of the IL4i1 reaction but remove any proteins greater than 3 kDa, including IL4i1 (∼78 kDa when glycosylated). Following the transfer of either filtered or unfiltered media ^51^, we observed that IL4i1^WT^ supernatants protected Kelly and SK-N-BE(2) cells from RSL3- or erastin-induced death (**Figure 3B, 3C, S2D and S2E**). By contrast, supernatants from IL4i1^ED^ transfected cells had no activity in suppressing ferroptosis. These experiments indicated that the enzymatic activity of IL4i1 is required for ferroptosis protection in bystander cells, rather than a non-enzymatic effect of IL4i1, such as binding and signaling via a cell surface receptor ^52,53^.

**Figure 3.**
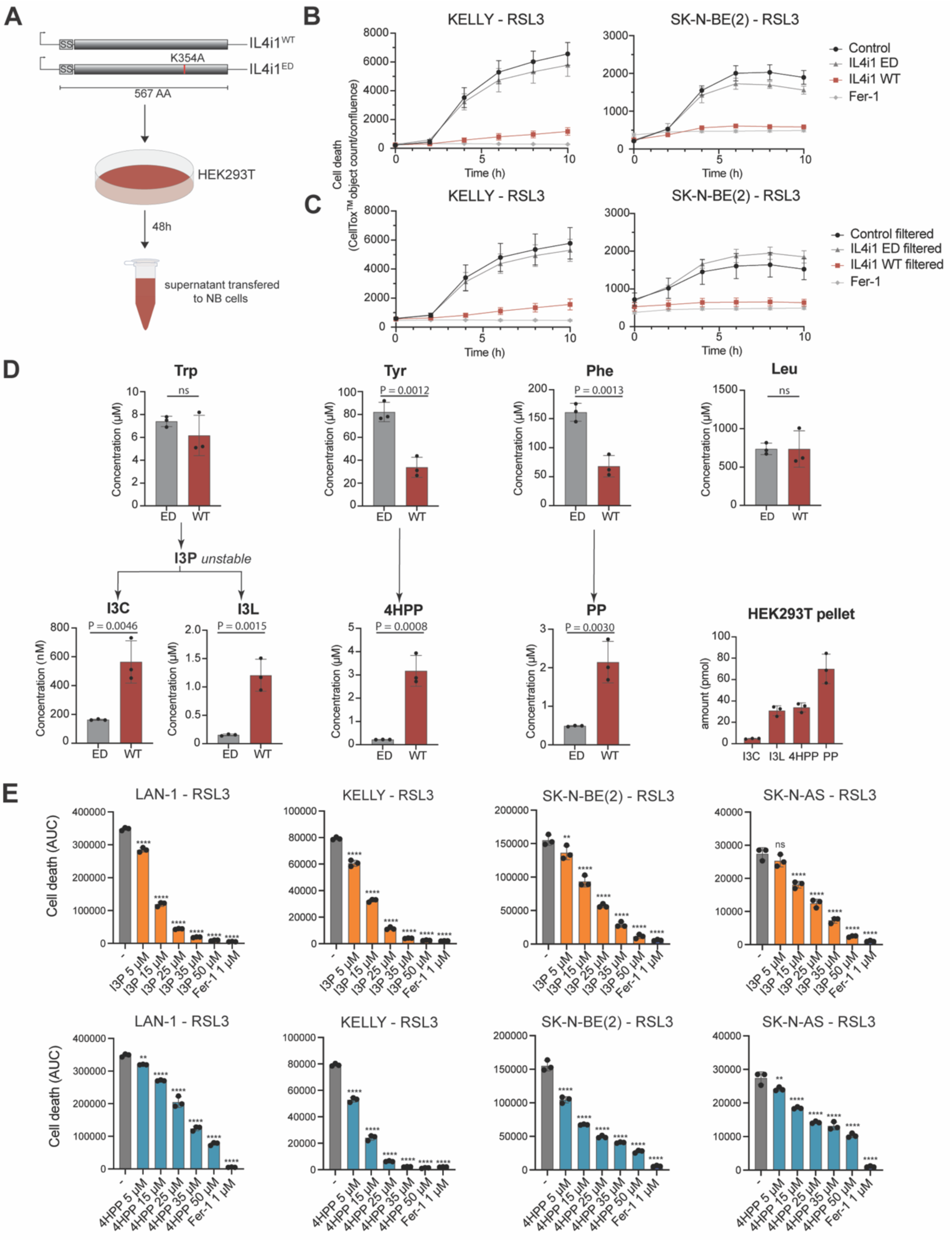
Analysis and effects of Il4i1 metabolism **(A)** Schematic of HEK293T cells transfected for 48 h with the human IL4i1 control (IL4i1^WT^) or enzyme-dead (IL4i1^ED^) cDNAs for supernatant transfer experiments. **(B)** Live-cell imaging of KELLY and SK-N-BE(2) cells that were treated for 24 h with supernatants collected from HEK293T cells. These HEK293T cells were either transfected with an empty vector (control) or transiently transfected with IL4i1^WT^ or enzyme-dead IL4i1^ED^ cDNAs. The NB cells were treated with 1 µM RSL3 and cell death was monitored over time using CellTox^TM^ Green. In the top graphs, the collected supernatants were used without further manipulation. **(C)** In the lower graphs, each supernatant was filtered to remove any material greater than 3 kDa. **(D)** Amino acid amounts (Trp, Tyr, Phe and Leu as a control) and corresponding metabolites amounts (I3C, I3L, 4HPP and PP) in the supernatants from HEK293T transfected cells with the human IL4i1 (IL4i1^WT^) or enzyme-dead (IL4i1^ED^) cDNAs for 48 h. The amounts of metabolites in the HEK293T cell pellets were also measured. Data are presented as mean ± s.e.m., n=3 samples. The P values were calculated using unpaired, one-tailed t test. ns: not significant. **(E)** Different human NB cell lines were challenged with RSL3 (0.1 µM for LAN-1, 0.25 µM for SK-N-BE(2), 0.5 µM for SK-N-AS and 1 µM for KELLY) in the presence or absence of titrated amount of I3P or 4HPP. 1 µM of Fer-1 was used as a control. Cell death was monitored over time with live cell imaging using CellTox^TM^ Green. Data are presented as the area under the curve (AUC) of a fixed time window during the transition from cell viability to death. Data are presented as mean ± s.e.m., n=3 samples. The experiment was replicated 3 times. One-way ANOVA with Dunnett’s multiple comparisons test was used after confirming normality. ns: not significant, **: P<0.01, ****: *P*< 0.0001.

Using targeted metabolomics, we analyzed the supernatants containing Il4i1^WT^ or IL4i1^ED^ for selected IL4i1 metabolites (**Figure 3D**). In the Il4i1^WT^ “milieu” we observed a slight decline in tryptophan concentration, which is used by HEK293T cells for growth and intermediary metabolism, and a more substantial reduction in tyrosine and phenylalanine compared to the IL4i1^ED^ milieu. These amino acid changes were accompanied by increased amounts of their cognate metabolites, including I3P-derived products, indole-3-carbaldehyde (I3C) and indole-3-lactate (I3L), as well as 4HPP and PP. Notably, the concentrations of these metabolites were lower than expected based on the correlations to the depletion of the source amino acid, possibly reflecting instability of certain metabolites and their degradation during sample extraction, or inaccurate response factors. Therefore, we next performed a titration of I3P and 4HPP using concentrations aligned with the levels of tryptophan and tyrosine the in serum. We tested the anti-ferroptotic activity of I3C and I3L and found that they do not protect NB cells from RSL3-or erastin-induced cell death (**Figure S2F**). We then proceeded with I3P and repeated the RSL3 and erastin challenge experiments across multiple human NB cell lines (**Figure 1C**). In each case, even at a concentration of 15 µM, I3P or 4HPP provided partial protection against ferroptosis (**Figure 3E and S2G**). Notably, IL4i1 simultaneously produces I3P and 4HPP at the same place and time, and together, they may have additive or synergistic effects. I3P is also rapidly imported by NB cells via yet to be determined transporters (**Figure S3A and S3B**), and may be further metabolized intracellularly. I3P is decomposed to other products ^34,54,55^, or potentially used for other metabolic pathways ^56^. Nevertheless, our results *in toto* suggest that naturally-secreted IL4i1 generated low micromolar concentrations of metabolites combinations that are sufficient to protect stressed high-risk NB cells from ferroptosis.

### IL4i1 suppresses NB death in 3D cultures

Expanding solid tumors contain necrotic cores deprived of oxygen and nutrients. These conditions can, in part, be mimicked in vitro using 3D tumor cultures. Cells in proximity to the core require stress-responsive pathways, including NRF2 and glutathione metabolism, for survival ^37^. We tested the contribution of IL4i1-derived metabolites to NB cell viability under 3D conditions by generating spheroids with LAN-1 cells in the presence or absence of I3P, 4HPP or Fer-1, and monitored cell death by live imaging with CellTox^TM^ Green. In untreated spheroids, green fluorescence was uniformly distributed, indicating widespread cell death. In contrast, spheroids treated with I3P, 4HPP or Fer-1 exhibited green fluorescence primarily in the core, while outer layers remained viable (**Figure 4A**). Treated spheroids also appeared more spherical, whereas untreated ones were irregular in shape. To substantiate these observations, we isolated individual spheroids and measured their ATP content as an indirect means of cell viability. Treated spheroids had approximately twice the ATP content as the controls (**Figure 4B**). Consistent with these findings, viability assays confirmed that titrated amounts of I3P or 4HPP improved spheroid cell survival relative to controls (**Figure 4C** and **4D**). While Fer-1 showed similar protective effects, Z-VAD-FMK and Necrostatin-1 failed to protect the cells, indicating that IL4i1 metabolites specifically target ferroptosis rather than other cell death mechanisms (**Figure 4E**). Notably, PP failed to promote cell viability, consistent with our previous findings ^33^.

**Figure 4.**
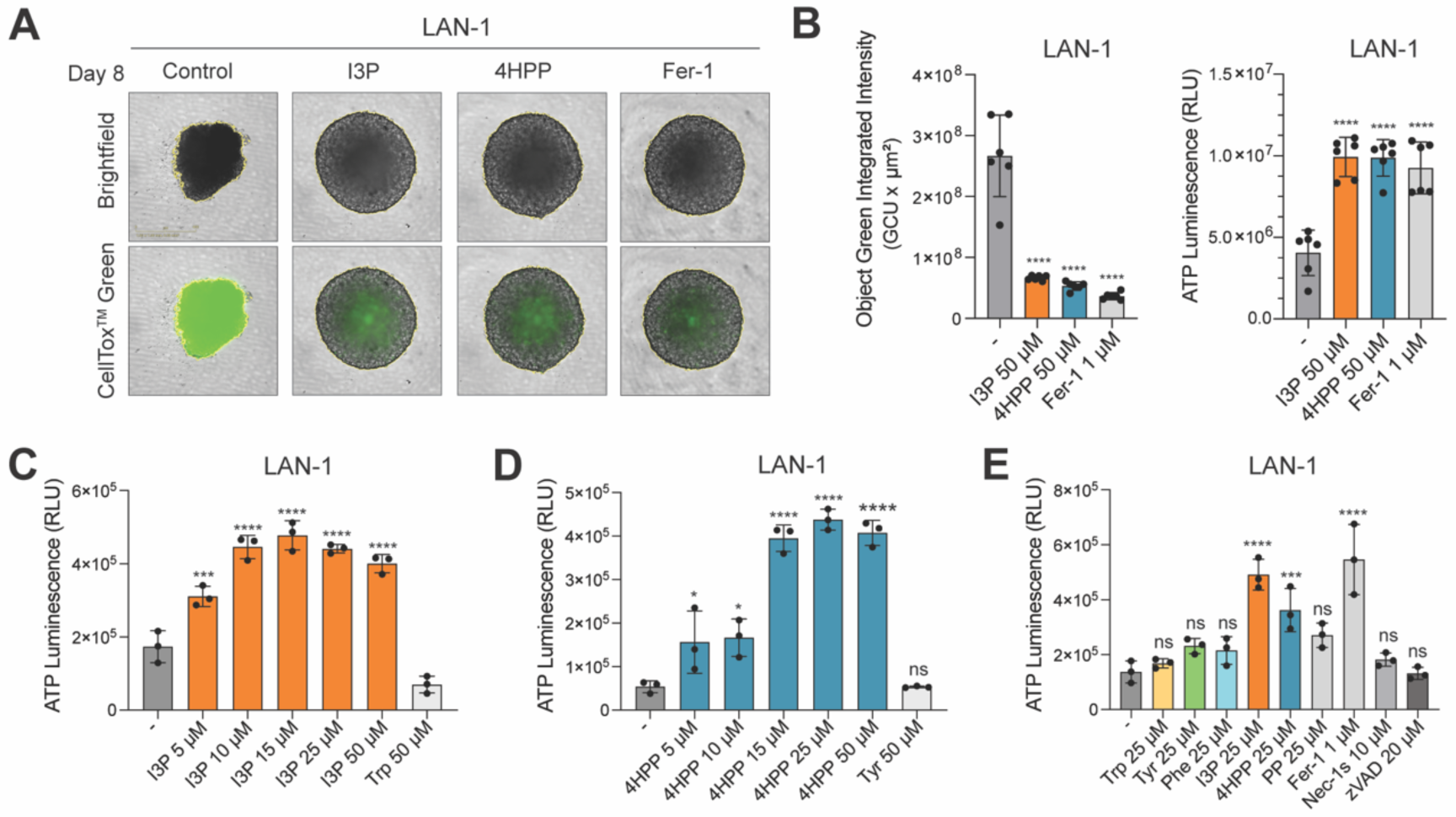
IL4i1 metabolites sustain viability in 3D **(A)** Representative images of LAN-1 cells grown in ultra-low attachment plates in the absence or presence of 50 µM I3P or 4HPP or 1 µM Fer-1 and stained with CellTox^TM^ Green. Media was added as a control. **(B)** Quantification of cell death within the spheroids using the green object intensity (CellTox^TM^ Green) or viability measuring the ATP luminescence (CellTiter-Glo® 3D). Data are presented as mean ± s.e.m., n=6 samples. The experiments were replicated 3 times. **(C, D)** Titrated amount of I3P and 4HPP were added to LAN-1 cells grown in ultra-low attachment plates. The cell viability was measured by ATP amounts (CellTiter-Glo® 3D). Data are presented as mean ± s.e.m. n=3 samples. The experiment was replicated 3 times. **(E)** LAN-1 cells were grown in 3D treated and with I3P, 4HPP or PP, their precursors (tryptophan, tyrosine or phenylalanine), Fer-1, Nec-1s or z-VAD and cell viability was quantified by ATP amounts (CellTiter-Glo® 3D). Data are presented as mean ± s.e.m. n=3 samples. One-way ANOVA with Dunnett’s multiple comparisons test was used in (B-E); ns: not significant, *:P<0.05, ***:P<0.001, ****:*P*< 0.0001.

### Mechanistic basis of IL4i1 suppression of NB ferroptosis

We next performed experiments to determine the mechanism(s) by which IL4i1 metabolites suppress ferroptosis in high-risk NB. We noted that multiple pathways are linked to I3P and 4HPP, including AHR activation, free radical scavenging, and NRF2 activation ^28^. Further, AHR has been linked to ferroptosis suppression ^57–59^. Conceivably, these pathways could influence ferroptosis outcomes either independently or in combination. Thus, we aimed to dissect the hierarchy of these pathways in the context of NB viability. We used RNA-seq to quantify gene expression in LAN-1 and SK-N-BE(2) cells following treatment with I3P or 4HPP over time. Unexpectedly, 4HPP elicited a robust NRF2 transcriptional signature compared to I3P, with an even more substantial effect observed after 24 hours (**Figure 5A, 5B and S4A**). This finding was corroborated by immunoblotting for NRF2 and selected NRF2 targets (SLC7A11, NQO1 and TXNRD1, **Figure 5C, 5D and S4B-D**). Based on these results, we generated NRF2-deficient (*NFE2L2*^-/-^) LAN-1 and KELLY cell lines (**Figure S4E and S4F**) and challenged them with RSL3 or erastin in the absence or presence of titrated doses of I3P or 4HPP. All high-risk NB cells depend on NRF2 to constitutively control their redox stress response. Accordingly, *NFE2L2*^-/-^ cells were exquisitely sensitive to RSL3 or erastin compared to their wild-type counterparts (**Figure 5E and S5H**). Nevertheless, I3P and 4HPP blocked ferroptosis in *NFE2L2*^-/-^ cells in a dose-dependent manner (**Figure 5F, S4G and S4H**). Additionally, spheroids derived from LAN-1 *NFE2L2*^-/-^ cells were also protected by I3P or 4HPP (**Figure 5G** and **5H**) compared to untreated spheroids. Given that I3P and 4HPP are free radical scavengers, we proceeded to evaluate their scavenging efficacy at different concentrations, as well as other downstream metabolites from I3P: I3C and I3L. While I3C and I3L exhibited no FRS activity (**Figure S4I**), both I3P and 4HPP demonstrated comparable scavenging effects to Fer-1, with I3P showing slightly greater activity than 4HPP (**Figure 5I**). Therefore, I3P and 4HPP modulate the NRF2 pathway coincident with scavenging free radicals, the latter of which may be sufficient to protect NRF2-deficient cells from ferroptosis.

**Figure 5.**
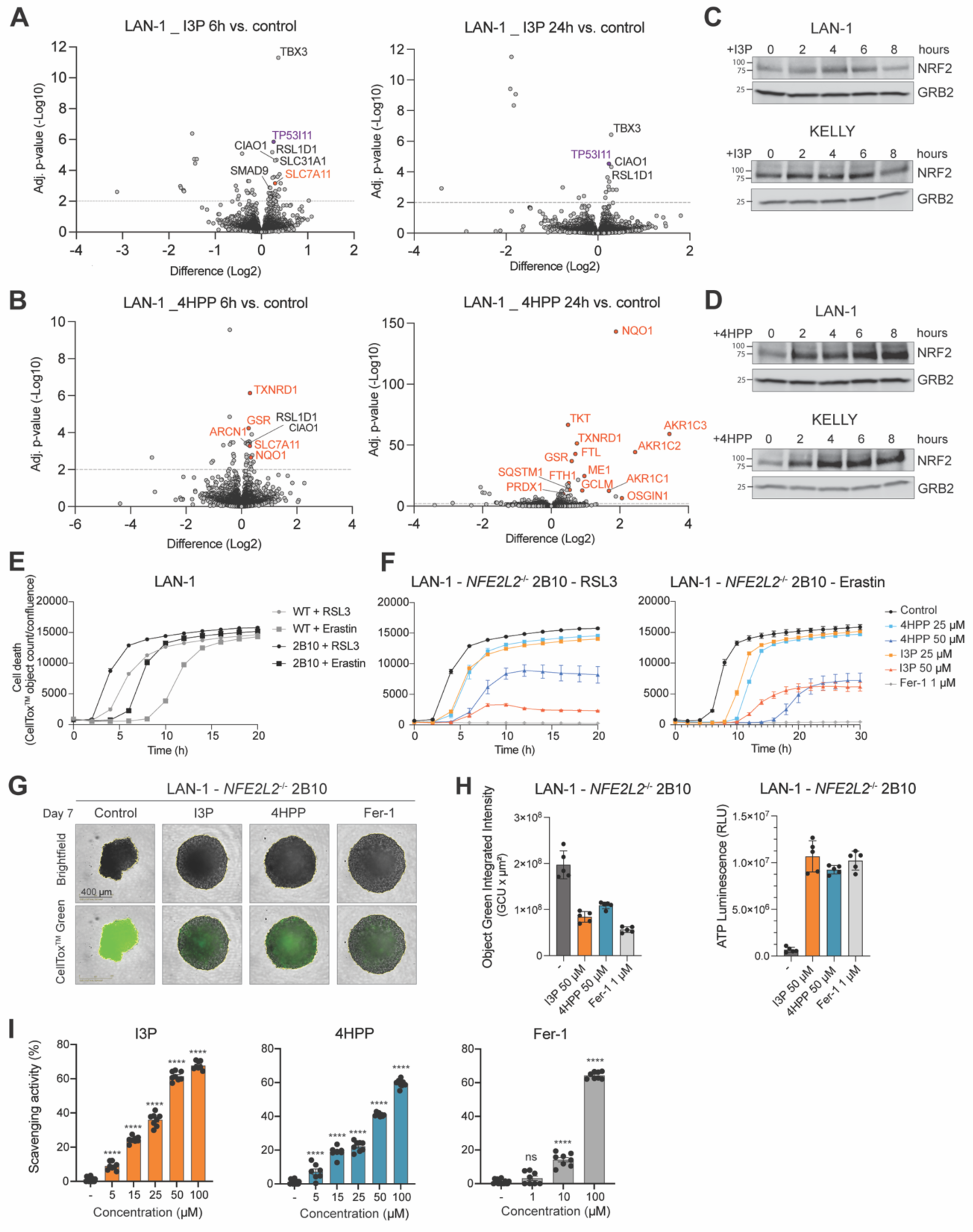
FRS and NRF2 activation are the key pathways required for IL4i1 metabolites to block ferroptosis **(A, B)** RNAseq analysis of LAN-1 cells treated with 50 µM I3P or 4HPP for 6 h and 24 h. Volcano plots compare untreated with treated cells. Key NRF2 (orange) and AHR (purple) target mRNAs are indicated. Averaged RNAseq data from three independent samples. **(C, D)** Immunoblotting analysis of NRF2 activation overtime in LAN-1 and KELLY cells following treatment with 50 µM I3P (C) or 50 µM 4HPP (D). **(E)** LAN-1 WT and NRF2-deficient LAN-1 2B10 clone were challenged with 0.1 µM RSL3 or 0.5 µM erastin. **(F)** NRF2-deficient LAN-1 2B10 clone was challenged with 0.1 µM RSL3 or 0.5 µM erastin in the absence or presence of I3P or 4HPP (25 µM and 50 µM) or Fer-1 (1 µM). Cell death was monitored over time by live cell imaging using CellTox^TM^ Green. Data are presented as mean ± s.e.m., n=3 samples. The experiment was replicated 3 times. **(G)** Representative images of NRF2-deficient LAN-1 2B10 clone grown in 3D in the absence or presence of 50 µM I3P, 4HPP or 1 µM Fer-1 and stained with CellTox^TM^ Green. Media was added as a control. **(H)** Quantification of cell death within the spheroids using the green object intensity (CellTox^TM^ Green) or viability measuring the ATP luminescence (CellTiter-Glo® 3D). Data are presented as mean ± s.e.m., n=5 samples. The experiments were replicated 3 times. **(I)** Cell-free scavenging activity of titrated concentrations of I3P, 4HPP or Fer-1 measured by the reduction in absorbance at 517 nm of the stable DPPH radical, relative an H20 control. Scavenging activity was calculated as: [(Control − Sample) / Control] × 100. Data represent the mean of 2 independent experiments, each with 4 technical replicates. Statistical analysis was performed using one-way ANOVA followed by Dunnett’s multiple comparisons test; ns: not significant, ****: P<0.0001.

As I3P is an AHR-activating metabolite ^34^, we quantified AHR activation over time following stimulation of LAN-1 and KELLY cells with I3P or 4HPP. While I3P rapidly induced AHR nuclear translocation, as previously described ^34^, 4HPP had no effect on AHR activation (**Figure S5A and S5B**). We further tested the relative contribution of AHR to ferroptosis resistance mediated by I3P and generated LAN-1 and KELLY cells lacking AHR (**Figure S5C and S5D**). *AHR*^-/-^ cells were protected from ferroptosis identically to the wild-type cells (**Figure S5E and S5F**). Thus, the AHR was dispensable for suppression of ferroptosis. These results implicate the FRS ability of I3P and 4HPP as a key component of their anti-ferroptotic properties. However, in “normal” high-risk NB cells with intact NRF2 signaling, the combined effects of NRF2 activation and FRS, independent of AHR activation, are used by the cells to enhance viability under stress.

### Loss of IL4i1 is protective in a mouse model of high-risk NB

We next tested the hypothesis that IL4i1 contributes to NB pathogenesis in vivo through a genetic approach. In the TH-MYCN; *Alk*^F1178L^ mice, NB formation is both penetrant and rapid, and the transcriptional landscape of the tumor cells is concordant with human high-risk adrenergic NB ^8^. We generated an *Il4i1*^-/-^ mouse strain, which showed a complete loss of IL4i1 protein (**Figure S6**). A complete knockout strategy was required here because different immune cells, including B and myeloid cell express IL4i1 in the mouse NB TME (**Figure 1B**). We measured tumor incidence and survival in 83 (see Methods for animal on and off-study criteria) TH-MYCN; *Alk*^F1178L^; *Il4i1*^-/-^ mice in comparison to 161 TH-MYCN; *Alk*^F1178L^ control mice. The IL4i1-deficient background was almost completely free of NB compared to the control TH-MYCN; *Alk*^F1178L^ animals (**Figure 6A** and **6B**). Interestingly, the small number of TH-MYCN; *Alk*^F1178L^; *Il4i1*^-/-^ mice that developed tumors were detected after 40 days on-study (i.e., after 4 imaging sessions) indicated that tumor formation, if it occurs, is also delayed relative to the control animals within the study window of 80 days. Therefore, our in vivo findings are consistent with the concept that IL4i1 is an immune-specific tumor promoting factor, generating pro-survival metabolites that act as oncometabolites.

**Figure 6.**
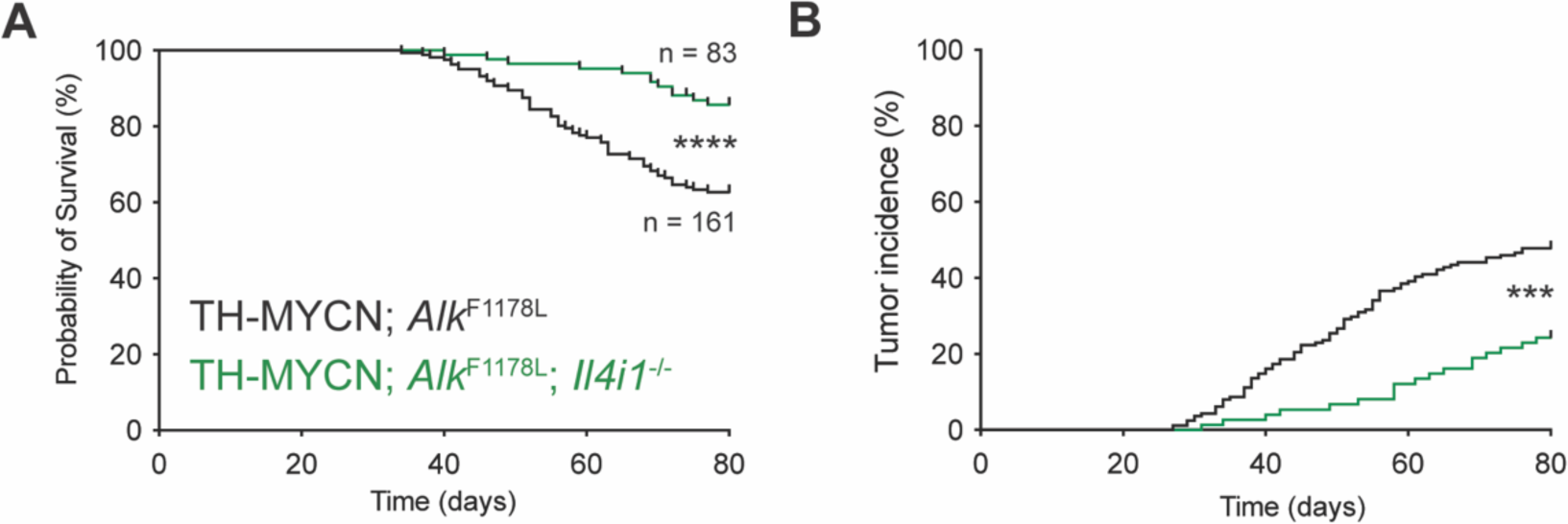
IL4i1 controls tumor formation and/or growth in a murine model of NB. **(A, B)** Probability of survival and tumor incidence of TH-MYCN; *Alk*^F1178L^; *Il4i1*^-/-^ mice relative to TH-MYCN; *Alk*^F1178L^ controls. TH-MYCN; *Alk*^F1178L^; *Il4i1*^-/-^ mice were imaged over the same ∼2-year period as the control animals ^8^. The Log-rank (Mantel-Cox) test was used as the primary statistic to estimate significance or non-significance; ***:*P*< 0.001, ****:*P*<0.0001.

## Discussion

IL4i1^+^ myeloid cells populate TMEs of all human solid tumors ^60,61^. Recently, IL4i1 mRNA expression has been used to define potentially pro-tumorigenic macrophages that infiltrate the TME ^62–65^, and IL4i1 is a hallmark of mregDCs ^66^, a population of activated dendritic cells that also invades tumors and tumor-draining lymph nodes. Two recent advances point to IL4i1 having pro-tumorigenic functions. First, metabolites generated by IL4i1-mediated oxidative deamination of tryptophan activate the AHR, which activates multiple pro-tumor pathways ^34^. Second, we discovered that I3P and 4HPP were ferroptosis suppressive molecules ^33^. As the TME itself and many chemotherapy agents trigger oxidative stress, malignant cells that survive hostile environments likely have increased requirements for adaptations that increase dependency on anti-oxidant pathways ^35–37^. Our results point to IL4i1 as a central cell-extrinsic, immune-derived pro-tumorigenic enzyme. First, I3P and 4HPP suppressed ferroptosis in human NB cell lines highly sensitive to perturbation of their oxidation stress defenses ^20–23^. Second, we found that the addition of IL4i1 metabolites also suppressed cell death in NB spheroids that manifest hypoxic cores. Third, our mechanistic experiments showed that the AHR was dispensable for ferroptosis protection, while NRF2 and FRS was required. These findings do not exclude other pro-tumorigenic pathways controlled by the AHR ^34^. Indeed, we imagine that IL4i1 could sustain tumor survival and proliferation by multiple parallel pathways; in our experiments, however, FRS and NRF2 activation are the core anti-ferroptotic pathways activated by IL4i1 metabolites. Fourth, our genetic experiments using a high-penetrance mouse model of human NB ^8,67^ provide important new evidence that IL4i1 mediates pro-tumor pathways in vivo as mice lacking IL4i1 were highly resistant to autochthonous oncogene-driven NB.

How do IL4i1-derived metabolites control ferroptosis? IL4i1 is secreted via the conventional endoplasmic reticulum pathway ^28^. Therefore, once in the extracellular space, the enzyme can oxidize its substrates (principally tryptophan, tyrosine and phenylalanine) to their keto forms. In previous work ^33^, and as shown herein for ferroptosis-sensitive NB cells, I3P (and likely 4HPP) is imported into cells via an unknown transport mechanism. The biochemical mechanism(s) of NRF2 activation by I3P and 4HPP remains unclear. Within the TME however, we propose that the metabolites of IL4i1 continuously supply local free radical scavenging and NRF2 activation. Therefore, increased IL4i1 expression and amounts may be positively correlated with the survival of malignant cells that have increased requirements for oxidative stress protection.

Our genetic approach clearly shows that the absence of IL4i1 is protective against the emergence of oncogene-driven NB. However, in this model system, it seems likely that the absence of IL4i1 affects an early step in tumor formation, as we did not observe TH-MYCN; *Alk*^F1178L^; *Il4i1*^-/-^ mice animals with tumors that subsequently regressed. Instead, few tumors were observed in the IL4i1-deficient background within the study window compared to controls. Recent information suggests tumor-infiltrating myeloid cells interact with transformed cells at the earliest steps in tumorigenesis and then help propel malignant growth, consistent with our previous assessment of TH-MYCN; *Alk*^F1178L^; *Ccr2*^-/-^ mice, where monocyte infiltration into tumors was reduced, an effect that partly protected the mice ^68^. While our results are consistent with this concept, defining and experimentally manipulating early events in cancer development remains challenging. One plausible interpretation of our genetic data is that initial oncogene-driven oxidative stress in transformed NB cells requires suppression from immune-sourced IL4i1. However, to test this idea, different model systems will be necessary, where tumors form to a point where their properties can be analyzed. Similarly, the key immune cell type necessary for IL4i1 production remains unclear in NB, or any other model: addressing this issue is intrinsically complex because IL4i1 is secreted into the milieu of the TME from multiple immune cell types. Based on our results reported here, and the widespread expression of IL4i1 in human and mouse myeloid cells ^60,63–66^ as well as other cells in tumor stroma ^46^, IL4i1 may be an attractive target for drug development and efficacy of an IL4i1 inhibitor such as an antibody, nanobody or irreversible small molecule may be facilitated be the fact that the enzyme works in the extracellular milieu.

### Limitations of the study

Our study has several limitations that require further experimentation. First, we used a complete IL4i1 loss-of-function approach for our in vivo experiments as multiple immune cells express IL4i1 in the NB TME. At first glance, the use of Cre-driven systems may be valuable in teasing apart which IL4i1^+^ cells are necessary for NB formation. However, this approach comes with a caveat: IL4i1 is a secreted protein and therefore elimination of IL4i1 in one immune cell type may not be sufficient to eliminate the effects of IL4i1 in the extracellular milieu that was generated from other cell type(s) within the TME. Further, we do not yet understand the amounts and dynamics of IL4i1 metabolite production in vivo. IL4i1 (if not regulated by an inhibitor or inactivation by, for example, extracellular proteases) likely generates I3P, 4HPP and PP continuously so long as tryptophan, tyrosine and phenylalanine are available. However, IL4i1-derived metabolites are imported by neighboring cells (probably in a differential way) ^33^ and can also be further metabolized to downstream products ^34^, both of which we also demonstrated here. Understanding the dynamic nature of this process will require a technology such as time- and cell-resolved tissue-level mass spectrometry to detect the metabolites, combined with a way to accurately measure the amounts of IL4i1 inside and outside different cell types. Third, while our experiments indicate NRF2 is a key pathway activated by IL4i1 metabolites in human NB cells, how they activate NRF2 remains unclear. Ascertaining the identity and hierarchy of the components of the NRF2 network activated by I3P and 4HPP may provide targets for drug development against oncogene-driven NB. Finally, we were not able to examine the changes in the phenotypes of immune and NB tumor cells in the absence of IL4i1, as tumor formation in TH-MYCN; *Alk*^F1178L^; *Il4i1*^-/-^ mice was too low. Therefore, different genetic systems (and tumor types) will be necessary to generate backgrounds where tumor formation occurs in an IL4i1-deficient background.

### Resource availability

All data are available from the corresponding authors upon request. The RNA-seq data can be found on GEO repository (GSE268972).

## Acknowledgments

We thank members of the PJM and PGT laboratories for assistance and feedback. We thank the members of the St. Jude Small Animal Imaging facility for their dedication to this project and Gustavo Palacios for metabolomics. Work in PJM’s laboratory is supported by FOR-2599 (type 2 tissue immunity), SFB-TRR 127 from the Deutsche Forschungsgemeinschaft (DFG), Boehringer Ingelheim and the Max-Planck-Gesellschaft. NBP is supported by NIH NCI F31 fellowship, award #: F31CA275320.

## Author contributions

C.G. and L. V. designed and performed research, analyzed and interpreted data and wrote the manuscript. L.K., J.R., L.Z. and A.B. designed and performed research, analyzed and interpreted data. N.P., B.S., E.K.A and A.Y analyzed and interpreted bioinformatic data; J.Y. provided key information and insight into the NB model along with in vitro experiments with NB cell lines. P.J.M. and P.G.T. designed the study, interpreted data, supervised research, and wrote the manuscript.

## Declaration of interests

The authors declare no competing interests that influenced the work herein.

## Supplemental Figures

**Figure S1.**
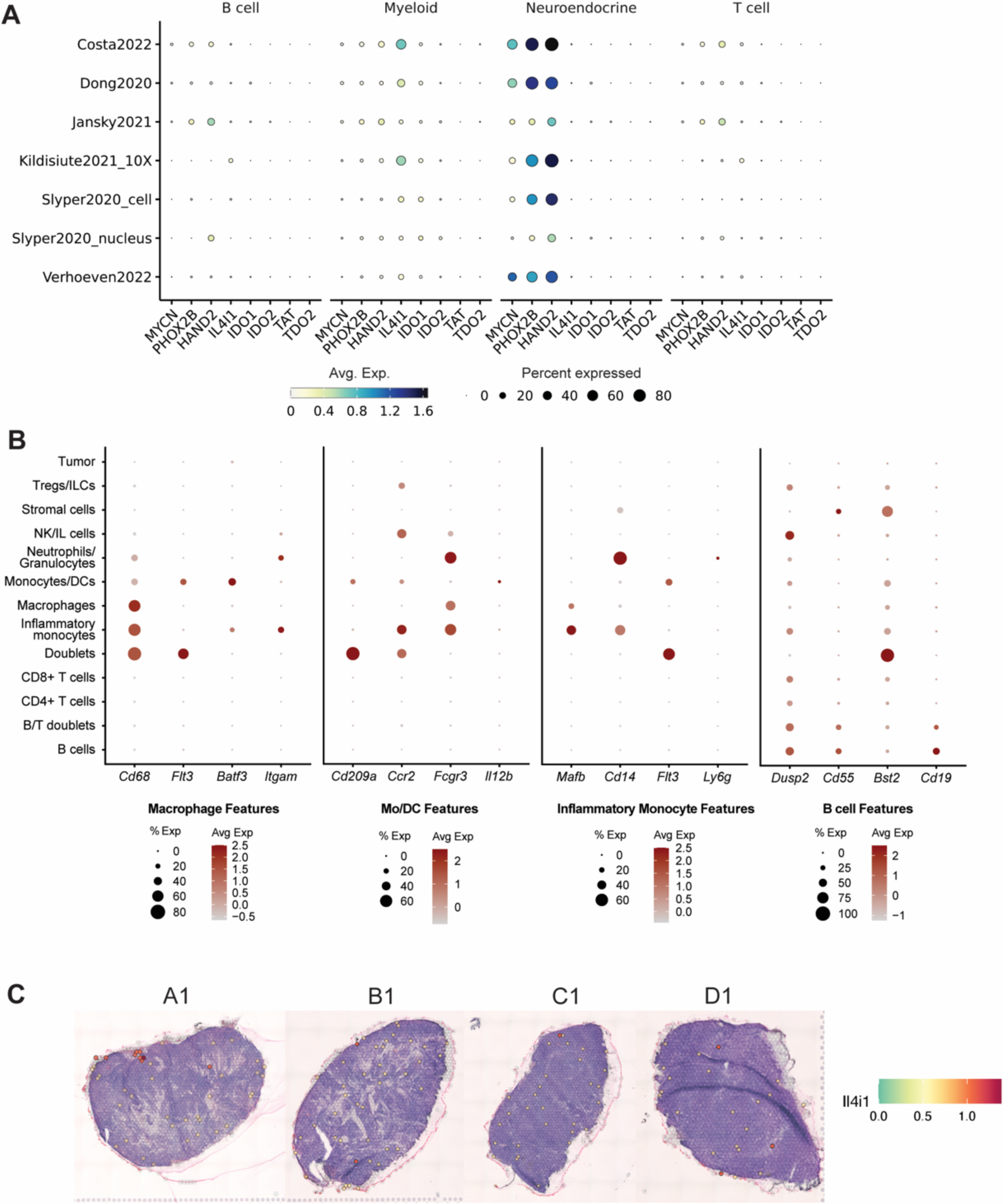
scRNAseq analysis of NB. Related to Figure 1. **(A)** Summary dot plots of human NB signature transcripts (MYCN, PHOX2B, HAND2) or aromatic amino acid metabolizing enzymes from seven scRNAseq studies. **(B)** TH-MYCN; *Alk*^F1178L^ Summary dot plots of featured transcripts from macrophages, Mo/DC, inflammatory monocytes and B cells from TH-MYCN; *Alk*^F1178L^ tumors. **(C)** Spatial transcriptomics analysis of IL4i1 mRNA from four individual TH-MYCN; *Alk*^F1178L^ tumors.

**Figure S2.**
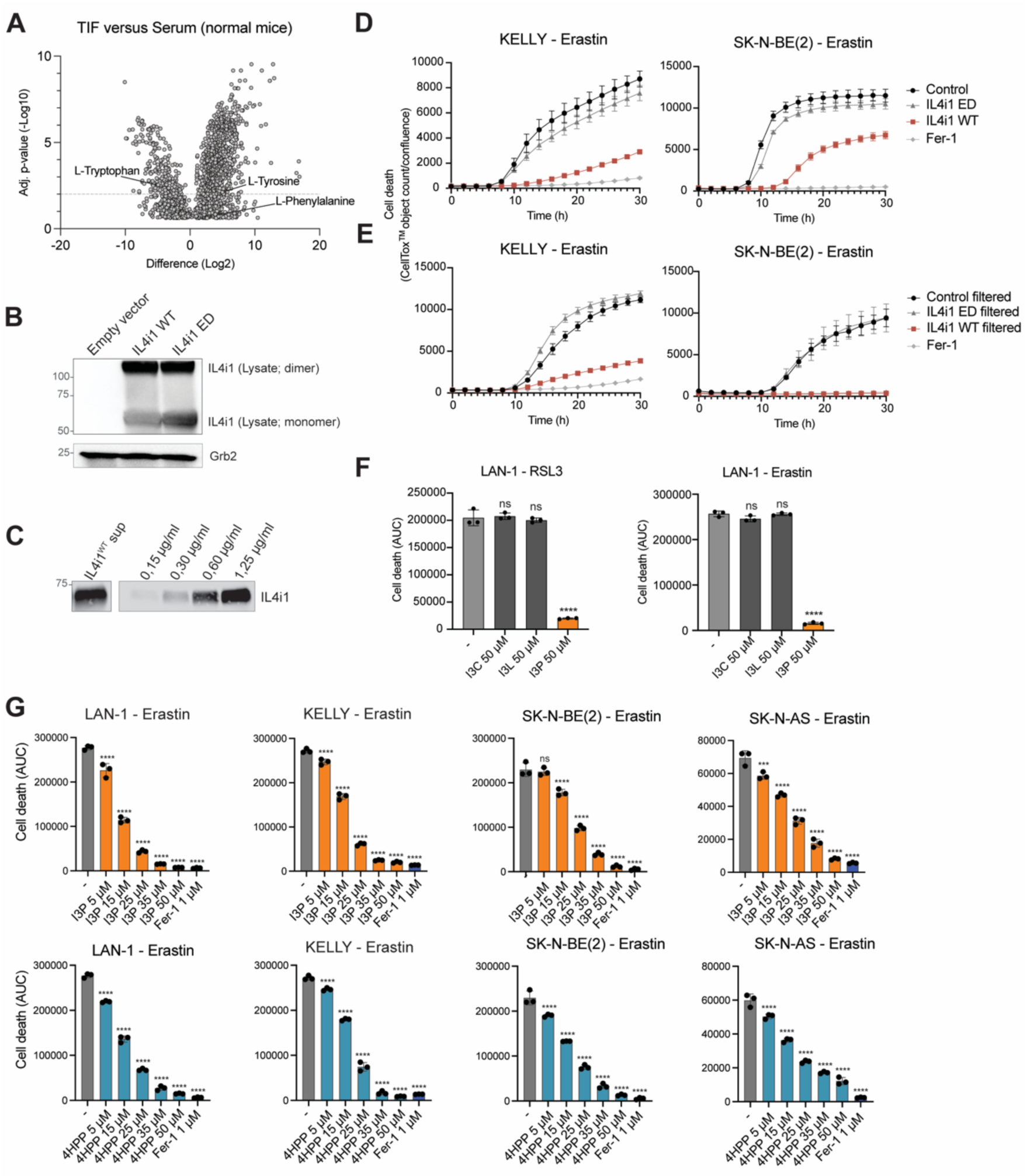
Generation of IL4i1 conditioned media for ferroptosis protection. Related to Figure 3. **(A)** Volcano plot depicting metabolite changes in tumors from TH-MYCN; AlkF1178L mice where fluid from the tumor was compared to serum from tumor free mice lacking the driver oncogenes but with the same genetic background. Mass data was collected in the positive mode from three tumors. **(B)** Immunoblotting analysis of transiently-transfected HEK293T cells. Cells were transfected with pcDNA3.1-Hygro as the empty vector control, or cDNAs (in pcDNA3.1-Hygro) encoding IL4i1 (IL4i1^WT^) or an enzyme-dead IL4i1 mutant (IL4i1^ED^) (see Figure 3A). Lysates were probed with an antibody specific for human IL4i1. Note that IL4i1 is secreted via the conventional ER pathway. When overexpressed, monomers and dimers are detected in the total lysate as indicated. **(C)** Immunoblotting of IL4i1^WT^ from transiently-transfected HEK293T cells compared to titrated amount or recombinant human IL4i1-C-Strep-tag. **(D, E)** Live-cell imaging of KELLY and SK-N-BE(2) cells that were treated for 24 h with supernatants collected from HEK293T cells. These HEK293T cells were either transfected with an empty vector (control) or transiently transfected with IL4i1^WT^ or enzyme-dead IL4i1^ED^ cDNAs. The NB cells were treated with 5 µM erastin and cell death was monitored over time with live cell imaging using CellTox^TM^ Green. In the top graphs, the collected supernatants were used without further manipulation. **(E)** In the lower graphs, each supernatant was filtered to remove any material greater than 3 kDa. **(F)** LAN-1 cells were challenged with 0.1 µM RSL3 or 0.5 µM erastin in the absence or presence of I3C, I3L or I3P (50 µM) and cell death was monitored by live cell imaging. Data are presented as the area under the curve (AUC) of a fixed time window during the transition from cell viability to death. Data are presented as mean ± s.e.m., n=3 samples. The experiment was replicated 3 times. **(G)** Different NB cell lines were challenged with erastin (0.5 µM for LAN-1 and SK-N-BE(2), 2.5 µM for SK-N-AS and 5 µM for KELLY) in the presence or absence of titrated amount of I3P or 4HPP. 1 µM Fer-1 was used as a control. Cell death was monitored over time with live cell imaging using CellTox^TM^ Green. Data are presented as the area under the curve (AUC) of a fixed time window during the transition from cell viability to death. Data are presented as mean ± s.e.m., n=3 samples. The experiment was replicated 3 times. One-way ANOVA with Dunnett’s multiple comparisons test was used for (F) and (G) after confirming normality. ns: not significant, **: P<0.01, ***:P<0.001, ****: *P*< 0.0001.

**Figure S3.**
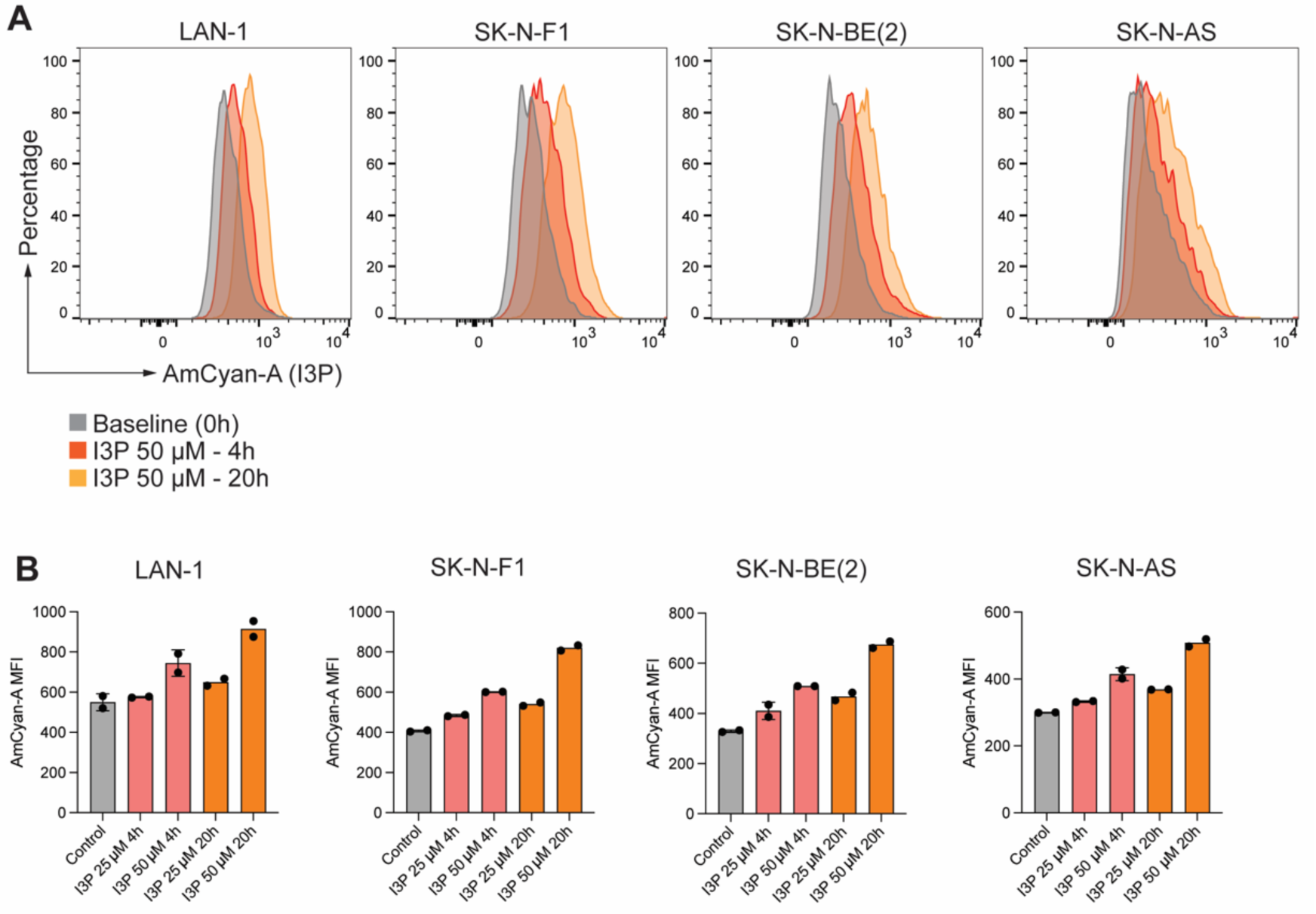
I3P is imported into NB cells **(A)** Histograms showing the uptake of I3P as measured by AmCyan-A fluorescence in different NB cell lines after 4h or 20h. **(B)** Mean fluorescence intensity (MFI) following I3P titrations in different NB cell lines after 4h and 20h quantified by flow cytometry of two independent samples.

**Figure S4.**
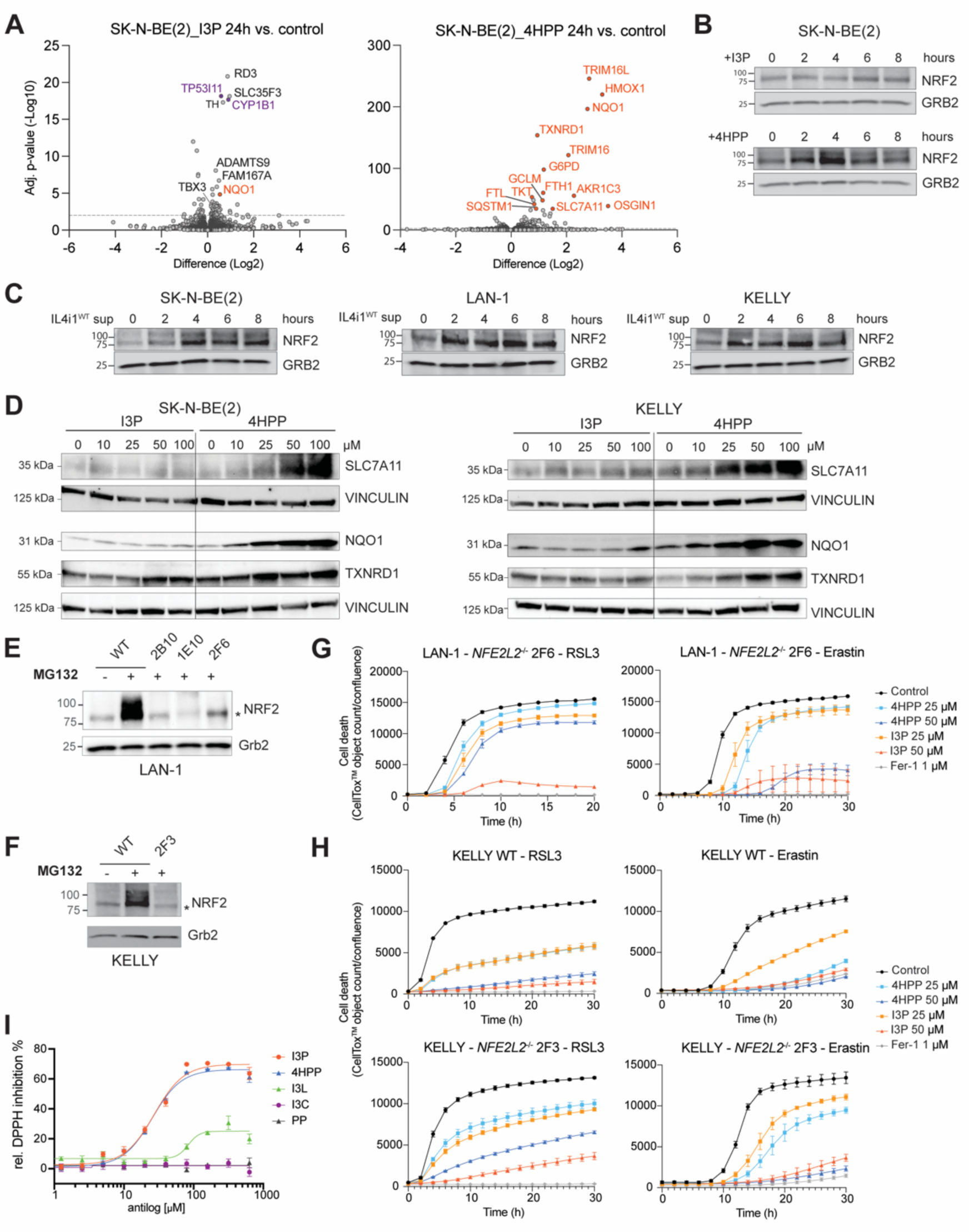
NRF2 response to IL4i1 metabolites. Related to Figure 5. **(A)** RNAseq analysis of SK-N-BE(2) cells treated with 50 µM I3P or 4HPP for 24 h. Volcano plots compare untreated with treated cells. Key NRF2 (orange) and AHR (purple) target mRNAs are indicated. Averaged RNAseq data from three independent samples. **(B)** Immunoblotting analysis of NRF2 activation overtime in SK-N-BE(2) cells following treatment with 50 µM I3P or 4HPP. **(C)** Immunoblotting analysis of NRF2 activation overtime in SK-N-BE(2), KELLY and LAN-1 cells following treatment with supernatant collected from HEK293T cells that were transiently transfected with control (IL4i1^WT^) for 48 h. **(D)** immunoblotting analysis of NRF2 target genes in SK-N-BE(2) and KELLY cells following stimulation with titrated amounts of I3P or 4HPP. Stimulations were performed for 24 h (SLC7A11) or 48 h (NQO1, TXNRD1) and each time point was compared to a corresponding Vinculin loading control. The experiment was replicated 3 times. **(E)** Analysis of NRF2-deficient LAN-1 clones. MG132 (20 µM) was added for 2 h to inhibit proteosome function and allow accrual of NRF2. *Indicates a background band. **(F)** Analysis of NRF2-deficient KELLY clones. MG132 (5 µM) was added for 6 h to inhibit proteosome function and allow accrual of NRF2. *Indicates a background band. **(G)** Analysis of cell death by Incucyte imaging in an additional NRF2-knockout clone as compared to Figure 5F. Data are presented as mean ± s.e.m., n=3 samples. The experiment was replicated 3 times. **(H)** KELLY WT cells and NRF2-deficient KELLY 2F3 clone were challenged with 1 µM RSL3 or 5 µM erastin in the absence or presence of I3P or 4HPP (25 µM and 50 µM) or Fer-1 (1 µM) and cell death was monitored over time by live cell imaging using CellTox^TM^ Green. Data are presented as mean ± s.e.m., n=3 samples. The experiment was replicated 3 times. **(I)** Radical scavenging activity of the listed metabolites was measured by quantifying the inhibition of the free radical DPPH (% absorbance) across increasing metabolite concentrations shown in antilog [µM]. Representative data are shown as mean ± SEM of 4 replicates.

**Figure S5.**
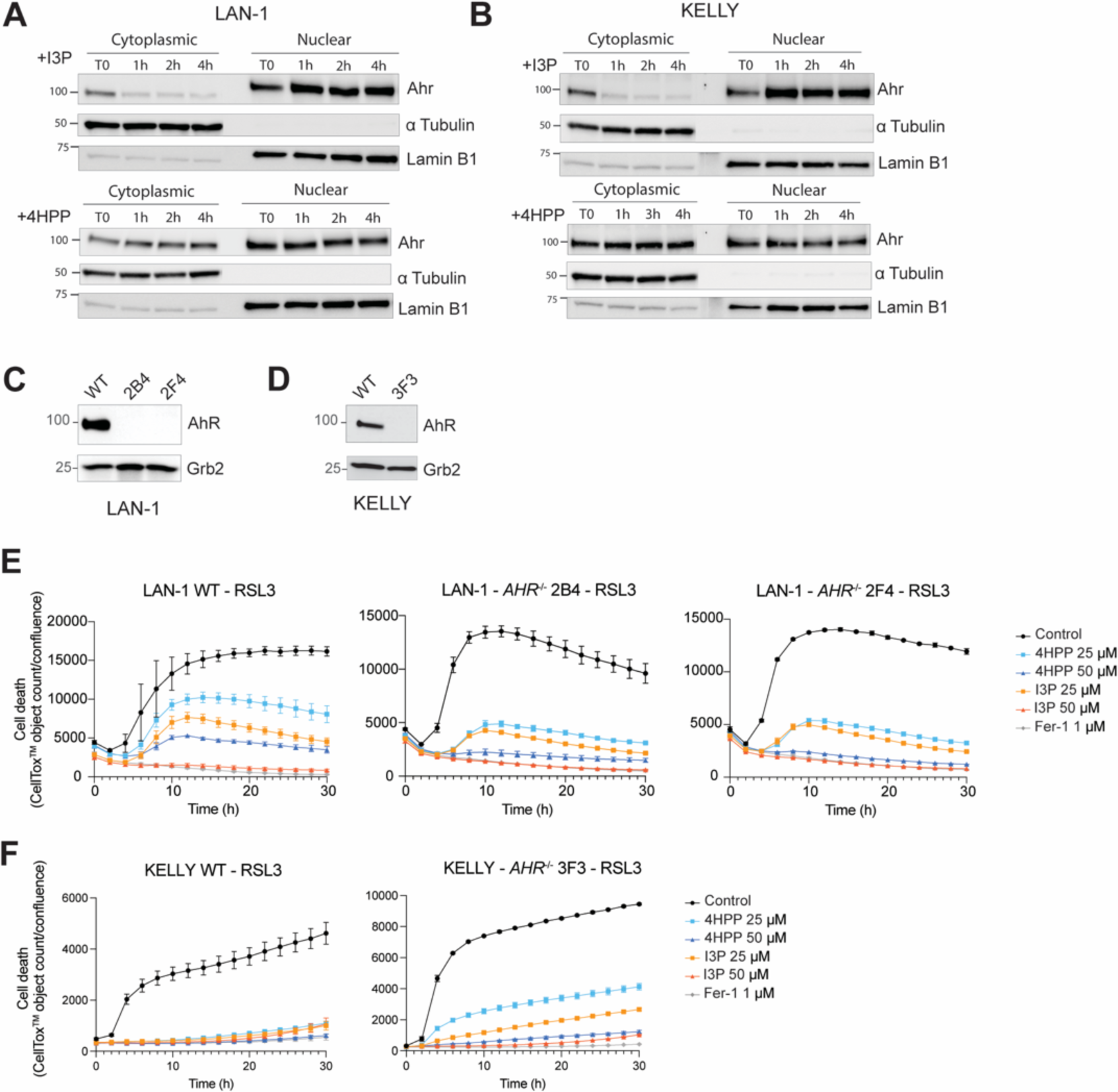
Role of AHR in anti-ferroptosis signaling. Related to Figure 5. **(A, B)** Immunoblotting analysis of AHR activation overtime in LAN-1 and KELLY cells following treatment with 50 µM I3P or 4HPP. Proteins were isolated from nuclear and cytosolic fractions. α Tubulin and Lamin B1 are cytosolic and nuclear markers, respectively. **(C, D)** Immunoblotting analysis of AHR-deficient LAN-1and KELLY clones. **(E)** WT and AHR-deficient LAN-1 NB cells were challenged with 0.1 µM RSL3 in the absence or presence of I3P or 4HPP (50 µM) or Fer-1 (1 µM) and cell death monitored over time by live cell imaging using CellTox^TM^ Green. Data are presented as mean ± s.e.m., n=3 samples. The experiment was replicated 3 times. **(F)** WT and AHR-deficient KELLY cells were challenged with 1 µM RSL3 in the absence or presence of I3P or 4HPP (50 µM) or Fer-1 (1 µM) and cell death monitored over time by live cell imaging using CellTox^TM^ Green. Data are presented as mean ± s.e.m., n=3 samples. The experiment was replicated 3 times.

**Figure S6.**
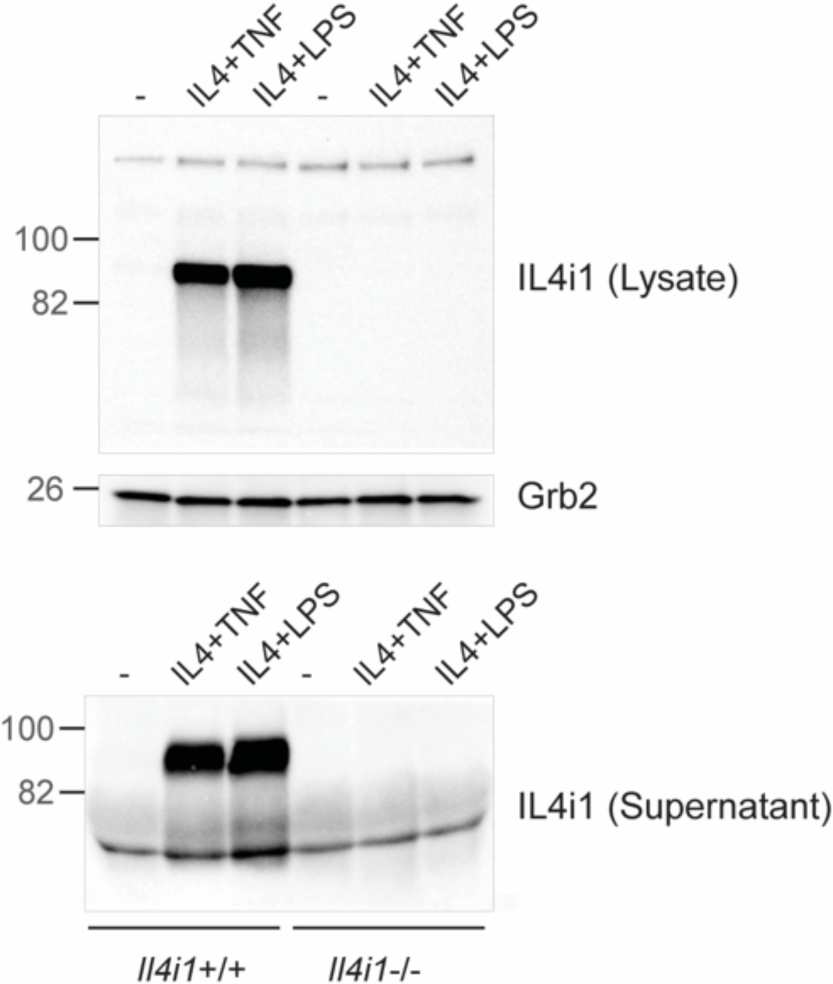
Antibody verification of IL4i1-deficient mice. Related to Figure 6. CSF-1 bone marrow-derived macrophages from control or *Il4i1*^-/-^ mice were unstimulated (-) or stimulated with IL4+TNF or IL4+LPS for 24 h and the cell lysate or cell supernatant probed for murine IL4i1 with a polyclonal antibody.

## Methods

### Visualization and analysis of the NBAtlas Resource

Harmonized single-cell and single-nucleus RNA sequencing data from six human neuroblastoma datasets were obtained from the publicly available NBAtlas resource (DOI: 10.17632/yhcf6787yp.2).

The processed Seurat object (seuratObj_NBAtlas_share_v20240130.rds) was loaded into R (v4.5.0) and used for all downstream analysis and visualization. Figures were generated using standard functions from Seurat (v5.3.0) and SCpubr (v2.0.2). Code is available at https://github.com/nbpeterson3/il4i1-viz.

### Cell lines

The human neuroblastoma cells lines LAN-1, SK-N-BE(2), KELLY, SK-N-AS and SK-N-F1 were cultured in DMEM-F12 (Gibco) supplemented with 10% FBS (Gibco) and 1% penicillin-streptomycin (Lonza). Human embryonic kidney 293T (HEK293T, CVCL_0063) cells were cultured in DMEM (Gibco) with 10% FBS (Gibco) and 1% penicillin-streptomycin (Lonza). All cell lines were cultured at 37°C in a humidified incubator under 5% CO2. All cell lines were tested for mycoplasma contamination using MycoStrip™ detection kit (Invivogen) upon initial thawing and then further tested every 3-4 weeks. Cell line identity was verified using short tandem repeat profiling service from Eurofins Genomics (Ebersberg, Germany).

### Neuroblastoma mouse model

Mouse NB experiments including study enrollment and off-study criteria (described in detail below) were performed at St. Jude Children’s Research Hospital as previously described ^8^ and approved by the internal Institutional Animal Care and Use Committee. Mouse breeding and housing at the Max-Planck-Institute of Biochemistry was approved by the institutional animal welfare officer according to animal licenses approved by the government of Upper Bavaria (Die Regierung von Oberbayern).

### Mouse models and quantification of autochthonous tumor growth

IL4i1-deficient mice were generated from ES cells supplied from a Lexicon gene trap ES line from MMMRC (B6;129S5-*Il4i1*^tm1Lex/Mmucd^). TH-MYCN; *Alk*^F1178L^ mice have been described previously ^8^. Mice were maintained by interbreeding to minimize space and the numbers of mice used. The genetic backgrounds of all mice used was mixed and originated from the permissive 129v/J background (TH-MYCN) ^10,11^ and C57BL/6. Following genotyping, mice were gender-segregated and assigned to imaging groups. For ultrasound imaging, fur was removed from the ventral side of each animal using Nair. Ultrasound imaging was performed all currently on-study enrolled mice weekly St. Jude Center for In Vivo Imaging and Therapeutics using a VEVO-2100, and determined tumor volumes using VevoLAB 3.0.0 software. All ultrasound data were initially acquired in a blinded fashion by the staff of the St. Jude Children’s Research Hospital Small Animal Imaging Core.

### Murine neuroblastoma study inclusion and exclusion criteria

Inclusion criteria have been described in detail before ^8^. Briefly, these include (i) presence of the TH-MYCN allele, (ii) heterozygosity for one *Alk* mutant allele and (iii) an age range from 1-3 weeks post-weaning. Either gender was used. Criteria for analysis were the presence or absence of an ultrasound-detectable tumor (quantified by staff unrelated to this study) at any point within a 7-week imaging period, where mice were imaged once per week. Off-study endpoints were (i) an animal welfare issue warranting euthanasia at any point, (ii) a tumor of >500 mm^3^, (iii) a tumor whose growth rate indicates a size of >500m^3^ would be reached before the next imaging session, (iv) death, or (v) 7 consecutive imaging sessions. For final analysis and survival data, the age of all mice was averaged to 80 days to account for minor variations in days of age at the first imaging session and survival and incidence analysis was performed in Prism.

### Mouse housing and diet

Mice at St. Jude Children’s Research Hospital were bred and housed within a single cubicle within a larger room. Mice were maintained in a 12 h day-night cycle, with constant temperature and humidity. Mice at the Max-Planck-Institute of Biochemistry were housed in open cages in a large room with a capacity for 200-250 cages and maintained on a 14-10 h day-night cycle in accordance with requirements from the animal facility. All mice were fed ad libitum using food approved by each respective animal facility.

### Cell transfection

HEK293T cells were seeded in a 6-well plate at a density of 0.5 x 10^6^ cells per well. After 24h they were transiently transfected with plasmids containing cDNA encoding WT IL4i1 or the enzyme-dead K354A mutant using Lipofectamine 3000 (Invitrogen) according to the manufacturer’s instructions. A mock transfection with an empty vector (pcDNA3.1-Hygro) was done as a control. After 6 h, the medium was refreshed, and the supernatant was collected 48 h later.

### Live cell imaging and quantification of cell death

Cells were seeded in a 96-well plate (in the presence of 1:6000 CellTox^TM^ Green and treated with the indicated concentrations of the compounds. After 24 h, ferroptosis was induced using erastin or RSL3 at different concentrations depending on the cell line. Cell death was monitored overtime using the IncuCyte S3 live-cell imaging device. Scans were acquired every 2 h for 30 h with five images per replicate using a 10x objective.

### 3D cell culture

Cells were seeded in a 96-well ultra-low attachment (ULA) microplate (S-BIO) using 800 cells/well in the presence of 1:6000 CellTox^TM^ Green and treated with media or compounds (50 µM of I3P or 4HPP or 1 µM of Fer-1). The plate was then centrifuged for 10 min at 300 × g. Half of the medium was refreshed every 2 days, avoiding disturbance of the developing spheroid. Cell death within the spheroid was monitored using IncuCyte S3 live-cell imaging and CellTox^TM^ Green (Promega). To evaluate cell viability based on ATP content within the spheroids, CellTiter-Glo® 3D (Promega) reagent was used, following the manufacturer’s protocol.

### Immunoblotting

Cells were washed with cold PBS and lysed in RIPA buffer containing a protease and phosphatase inhibitor cocktail (Thermo Scientific). Nuclear and cytoplasmic extracts were prepared using NE-PER™ extraction kit. Proteins were separated by SDS-polyacrylamide gel electrophoresis (4– 15% Criterion TGX Stain-Free protein gels, 5678085, Bio-Rad), transferred to nitrocellulose membranes (0.2 µm, 10600001, Amersham), and blocked in 3% milk in TBS containing 0.1% Tween-20 (Sigma). The membranes were incubated overnight at 4°C with the primary antibodies in 3% milk. Membranes were washed and incubated with 1:10,000 HRP-conjugated secondary antibodies Goat Anti-Rabbit and Goat Anti-Mouse before visualization with SuperSignal West Pico Substrate (34080; Pierce).

### Free radical scavenging activity assay

The stable free radical 1,1-diphenyl-2-picrylhydrazyl (DPPH) was used to determine the radical scavenging activity of the metabolites I3P, I3L, I3C, 4HPP and PP. The ferroptosis inhibitor Fer-1 was used as positive control. The metabolites were dissolved in PBS or PBS+DMSO. A 200 μM DPPH stock solution was prepared in methanol and pipetted in a 96-well plate. Equal volumes of each metabolite compound were added in quadruplicates for all the tested concentrations and mixed briefly with the DPPH. An equal volume of PBS or PBS+DMSO (respective to metabolite solvent) was used as control. After 10 min incubation, the absorbance was measured at 517 nm with a Tecan Spark microplate reader. The scavenging activity, leading to a decrease of DPPH absorbance, was calculated as relative absorbance to the respective control: (A517 control – A517 sample) / A517 control) x 100 %.

### CRISPR-Cas9–mediated gene knockout

To generate NRF2- and AHR-deficient NB cells, sgRNA sequences were designed using standard design tools (IDT) and cloned into the pX458 CRISPR vector (pSpCas9(BB)-2A-GFP, Addgene, #48138). LAN-1 and KELLY cells were seeded in a 6-well plate at 1 × 10^6^ cells per well. After 24 h, cells were transfected using Lipofectamine 3000 and the media was exchanged the next day. After 24 h, cells were sorted by FACS for GFP^+^ and single cells were added to 96-well plates containing 300 μL of media with 20% FBS. After expansion, verification of loss-of-function was done by immunoblotting and sequencing. The following sgRNA sequences were used:

*NFE2L2* sgRNA_1 (exon 2): GACAAGAACAACTCCAAA

*AHR* sgRNA_2 (exon 2): GGCAGCAGGCTAGCCAAA

### Sample preparation for LC-MS/MS

HEK293T cells were transiently transfected with plasmids containing cDNA encoding WT IL4i1 or the enzyme-dead K354A mutant as previously described. After 48 h supernatant were harvested and immediately put on dry ice then stored at −80 °C. The pellets were washed with PBS and stored at −80 °C. Ice-cold extraction buffer (0.5 mL; ACN/MeOH/water, 2:2:1, v/v) was added to 50 µL of supernatant or to the cell pellets. The samples were vortexed thoroughly. Pellet samples were additionally subjected to ultrasonication for 10 minutes using a Bioruptor (Diagenode), with 10 cycles of 30 seconds at high intensity followed by 30-second pauses between each cycle. After centrifugation for 10 minutes at 12,000 × g at 8 °C, the resulting supernatants were collected and dried using a vacuum concentrator (SpeedVac).

### LC-MS/MS Data acquisition and analysis

Data acquisition was performed on a Q Exactive HF mass spectrometer (Thermo Scientific) coupled to a Vanquish Flex HPLC system (Thermo Fisher Scientific). Vacuum-dried metabolite extracts from supernatants were reconstituted in 200 µL of buffer A (0.1% formic acid), and 6 µL was injected for analysis. Extracts from pellets were reconstituted in 40 µL of buffer A, and 3 µL was injected. Metabolite separation was carried out on a Kinetex F5 column (2.1 × 100 mm, 2.6 µm; Phenomenex) at a flow rate of 200 µL/min. The gradient was as follows: buffer B (100% ACN, 0.1% formic acid) was held at 0% for 2 minutes, linearly increased to 95% over 12 minutes, maintained at 95% for 2 minutes, and then returned to 0% for a 4-minute re-equilibration. The mass spectrometer was operated in negative ion mode. Full MS1 scans were acquired from m/z 75 to 1000 at a resolution of 120,000. HESI source parameters were set as follows: sheath gas flow rate, 47 arbitrary units (AU); auxiliary gas flow rate, 10 AU; sweep gas flow rate, 2 AU; spray voltage, 3.20 kV; capillary temperature, 250 °C; and auxiliary gas heater temperature, 380 °C. Up to five top precursors were selected for MS2 fragmentation using stepped higher-energy collisional dissociation (HCD) with normalized collision energies of 20, 40, and 60. MS2 spectra were acquired at a resolution of 30,000. The automatic gain control (AGC) target values were 3 × 10⁶ for MS1 and 1 × 10⁵ for MS2, with maximum injection times of 200 ms and 50 ms, respectively.

Peak areas corresponding to metabolites of interest were identified using MS1 high-resolution accurate mass, characteristic MS2 fragmentation patterns, and retention times, matched to those of unlabeled standards. Data processing was performed using Skyline software (version 25.1; MacCoss Lab, University of Washington). Concentrations were calculated based on previously established calibration curves generated from unlabeled standards.

### Sample preparation for RNA-seq

LAN-1 and SK-N-BE(2) cells were seeded in 6-well plates at a density of 0.4 × 10^6^ cells per well. After 2 days the cells were treated with 50 μM of I3P or 4HPP for 6 h or 24 h. The media was removed and cells were collected in TRIzol™ (Thermo Fisher Scientific) and stored at −80 °C for 1 week. 200 μL of chloroform was added to the samples and the RNA phase was collected after centrifuging for 10 min at 4°C at 12700rpm and added to 500 μL of 2-propanol. The samples were stored at −20 °C overnight. The supernatant was removed after centrifugation for 10 min at 4°C at 12700rpm and 1ml of EtOH 70% was added to the RNA pellet. The supernatant was removed after 5 min centrifugation and the RNA dried for 5 min at RT. 25 μL of pure distilled water was then added.

### NGS data set analysis

After quality-checking using the tool FastQC (v.0.11.7) (http://www.bioinformatics.babraham.ac.uk/projects/fastqc), the files were mapped to the human genome (Genome build *GRCh38*) downloaded from Ensembl using the star aligner (v. 2.7.10b) ^69^ using default parameters. The mapped files were then quantified on a gene level based on the ensembl annotations, using the featureCounts (v. 2.0.4) tool from the SubRead package ^70,71^, discarding multi-overlapping reads. Using the DESeq2 package (R 4.3.2, DESeq2 version 1.42)^72^ the count data was normalized by the size factor to estimate the effective library size. A filtering step for removing genes with less than 10 reads in at least three samples was used. Genes with an adjusted p-value of <= 0.01 were than considered to be differentially expressed for downstream analysis. After calculating the gene dispersion across all samples, the comparison of the two different conditions resulted in a list of differentially expressed genes.

### Quantification and statistical analysis

All mice included into the study were analyzed sequentially. No power analysis was performed before the study began because the rate of tumor formation in the mutant ALK backgrounds in the St. Jude vivarium was unknown at the initiation of the study. Therefore, measurement of tumor formation was longitudinal with corresponding analysis to estimate significance or non-significance in tumor formation over time. At the cessation of the study (80 days, per requirement of the animal protocol), all primary data was independently checked against the original database by two people and entered into GraphPad Prism v7 for survival curve comparison. The Log-rank (Mantel-Cox) test was used as the primary statistic to estimate significance or non-significance. Two-tailed *t* tests or Kruskal-Wallis tests corrected for multiple comparisons were used as appropriate. For comparison of two groups, after confirming normality, data were analyzed with two-tailed unpaired Student’s *t* test; if data did not follow normal distribution, the non-parametric Mann-Whitney U-test was used. When comparing more than two groups and after confirming normality, one-way ANOVA followed by Dunnett’s multiple comparisons test was used. Two-way repeated measures ANOVA and Sidak’s multiple-comparisons test was used to analyze data in repeated-measures designs. All statistical analyses were performed using GraphPad Prism software (version 10.2.0, GraphPad Software, San Diego, CA, USA). *P* values < 0.05 were considered to be statistically significant.

## Notes

### Competing Interest Statement

The authors have declared no competing interest.

